# Experimental co-transmission of simian immunodeficiency virus (SIV) and the macaque homologs of the Kaposi sarcoma-associated herpesvirus (KSHV) and Epstein-Barr virus (EBV)

**DOI:** 10.1101/432351

**Authors:** A. Gregory Bruce, Serge Barcy, Jeannette Staheli, Helle Bielefeldt-Ohmann, Minako Ikoma, Kellie Howard, Timothy M. Rose

**Affiliations:** Center for Global Infectious Disease Research, Seattle Children’s Research Institute, Seattle, Washington, United States of America; Department of Pathobiology, University of Washington, Seattle, Washington, United States of America; Department of Pediatrics, University of Washington, Seattle, Washington, United States of America; Washington National Primate Research Center, University of Washington, Seattle, Washington, United States of America

## Abstract

Macaque RFHV and LCV are close homologs of human KSHV and EBV, respectively. No experimental model of RFHV has been developed due to the lack of a source of culturable infectious virus. Screening of macaques at the Washington National Primate Research Center detected RFHV in saliva of SIV-infected macaques from previous vaccine studies. A pilot experimental infection of two naïve juvenile pig-tailed macaques was initiated by inoculation of saliva from SIV-infected pig-tailed and cynomolgus macaque donors, which contained high levels of DNA of the respective species-specific RFHV strain. Both juvenile recipients developed SIV and RFHV infections with RFHV DNA detected transiently in saliva and/or PBMC around week 16 post-infection. One juvenile macaque was infected with the homologous RFHVMn from whole saliva of a pig-tailed donor, which had been inoculated into the cheek pouch. This animal became immunosuppressed, developing simian AIDS and was euthanized 23 weeks after inoculation. The levels of RFHV DNA in saliva and PBMC remained below the level of detection after week 17, showing no reactivation of the RFHVMn infection during the rapid development of AIDS. The other juvenile macaque was infected with the heterologous RFHVMf from i.v. inoculation of purified virions from saliva of a cynomolgus donor. The juvenile recipient remained immunocompetent, developing high levels of persistent anti-RFHV and -SIV antibodies. After the initial presence of RFHVMf DNA in saliva and PBMC decreased to undetectable levels by week 19, all attempts to reactivate the infection through additional inoculations, experimental infection with purified SRV-2 or SIV, or immunosuppressive treatments with cyclosporine or dexamethasone were unsuccessful. An heterologous LCV transmission was also detected in this recipient, characterized by continual high levels of LCVMf DNA from the cynomolgus donor in both saliva and PBMC, coupled with high levels of anti-LCV antibodies. The macaque was sacrificed 209 weeks after the initial inoculation. Low levels of LCVMf DNA were detected in salivary glands, tonsils and other lymphoid organs, while RFHVMf DNA was below the level of detection. These results show successful co-transmission of RFHV and LCV from saliva and demonstrate differential lytic activation of the different gammaherpesvirus lineages due to presumed differences in biology and tropism and control by the host immune system. Although this initial pilot transmission study utilized only two macaques, it provides the first evidence for experimental transmission of the macaque homolog of KSHV, setting the stage for larger transmission studies to examine the differential activation of rhadinovirus and lymphocryptovirus infections and the pathological effects of immunosuppression.

## Introduction

Two members of the gammaherpesvirus subfamily, Kaposi’s sarcoma-associated herpesvirus (KSHV)/human herpesvirus 8 and Epstein-Barr virus (EBV)/human herpesvirus 4 infect humans and are associated with a number of malignancies and proliferative disorders. KSHV, genus Rhadinovirus (RV), is the etiologic agent of Kaposi’s sarcoma (KS), an endothelial cell-derived malignancy, and plays a role in the pathogenesis of several B-cell lymphoproliferative disorders, including multicentric Castleman disease (MCD) and primary effusion lymphoma (PEL) [1]. EBV, genus *Lymphocryptovirus* (LCV), is etiologically associated with lymphoproliferative B-cell disorders, including Burkitt’s and Hodgkin’s lymphomas, as well as epithelial-derived nasopharyngeal and gastric carcinomas [2]. The genomes of KSHV and EBV are in general co-linear, and the majority of genes show obvious sequence homology. However, each viral lineage is characterized by genetic insertions, duplications and mutations that profoundly differentiate the biology and pathology of the two viruses. In dually infected PEL cells, regulatory genes of both viruses can interact and suppress the lytic replication of each other [3–6]. In addition, both of these oncogenic viruses have evolved mechanisms to induce long-term viral latency by altering cellular gene expression using viral-encoded microRNAs (miRNAs). EBV and KSHV miRNAs target cellular pathways of apoptosis, cell-cycle control and immune-modulation, which enable the viral infections to persist [7–9].

We and others have shown that homologs of KSHV and EBV are present in macaques and other Old World primates, including chimpanzees, gorillas and mandrills [10–16]. Two lineages of rhadinoviruses related to KSHV have been identified in Old World primates [17] [18]. The RV1 rhadinovirus lineage consists of KSHV and closely related homologs in all Old World primate species examined. We identified the retroperitoneal fibromatosis herpesvirus (RFHV), the macaque RV1 homolog of KSHV, in KS-related tumors in macaques with simian AIDS at the Washington National Primate Research Center (WaNPRC). Distinct RFHV variants are present in each macaque species, including RFHVMn in pig-tailed macaques (*M. nemestrina*), RFHVMm in rhesus macaques (*M*. *mulatta*) and RFHVMf in cynomolgus macaques (*M. fascicularis*) [10, 19]. We have obtained the complete sequence of the genome of RFHVMn [20] and partial sequences of the RFHVMm and RFHVMf genomes [21–23]. The RFHV and KSHV genomes are highly conserved with strong sequence conservation. Both macaque and human RV1 rhadinoviruses encode a set of core genes conserved across the herpesvirus family, which function in virus replication and virion structure. The RV1 genomes also contain a group of conserved latency-associated genes, including the ORF73 latency-associated nuclear antigen (LANA) [24], which function to maintain latency, as well as a number of novel genes exhibiting sequence homology to cellular genes involved in mitosis, cell cycle regulation and immunity [20, 21, 25].

The RV2 rhadinovirus lineage consists of a set of closely related rhadinoviruses in different Old World primate species, including macaques, chimpanzees and gorillas [11, 14, 17, 18, 26]. No RV2 rhadinovirus has been detected in humans, suggesting that this viral lineage was lost during hominid evolution. The genomes of two variants of the RV2 rhadinovirus from rhesus macaques have been completely sequenced, RRV-17577 (Oregon National Primate Research Center) [27] and RRV26–95 (New England National Primate Research Center)[28]. We have sequenced the complete genome of the RV2 rhadinovirus from a pig-tailed macaque, MneRV2-J97167, at the WaNPRC [29] and have obtained partial sequences of MfaRV2, from a cynomolgus macaque [19, 22]. Overall, the genomes of MneRV2 and other macaque RV2 rhadinoviruses resemble those of the RV1 rhadinoviruses, although the RV2 rhadinoviruses lack a number of conserved RV1 genes implicated in immune evasion and pathogenesis and have an eight-fold amplification of an interferon regulatory gene.

Studies have shown that the RV1 and RV2 rhadinoviruses have different biological properties. RV1 rhadinoviruses show a strong tendency to develop latent infections [30–34], whereas the RV2 rhadinoviruses show a more lytic phenotype resulting in viral replication [22, 26, 35]. Members of the two rhadinovirus lineages utilize different cellular receptors [23, 36–42] suggesting different cellular tropisms with different infection outcomes.

Simian homologs of EBV, called lymphocryptoviruses (LCV), have also been detected in a wide variety of Old World primates, initially by cross-reactive antibodies and later by molecular methods [16]. The complete genome sequence has been determined for LCV from the rhesus macaque (*M. mulatta*) [43], herein designated MmuLCV,. The genome of MmuLCV was found to have a repertoire of viral genes identical to human EBV, with a high degree of amino acid sequence similarity, especially in the lytic infection proteins. The biology of MmuLCV infection in the rhesus macaque appears virtually identical to EBV infection in humans, with regard to cell tropism, persistence, cellular and humoral immunity, and secretion of infectious virus in saliva.

Little is known regarding the effects of KSHV and EBV co-infection in vivo. Since both viruses show an exclusive tropism for humans, animal models to study virus infection, persistence and pathogenesis have not been possible. However, models using mice reconstituted with human immune system components have been developed [44]. With such models, KSHV and EBV infections have been studied using oral inoculation or intraperitoneal injection of purified virus particles [45, 46]. These studies showed that KSHV and EBV establish infections in the human B-cell population in vivo. Subsequent studies showed that co-infection of the humanized mice resulted in an enhanced persistence of KSHV in co-infected EBV-transformed human B-cells. Increased lytic replication of EBV was detected in the KSHV-co-infected cells, which was associated with increased tumor formation [47]. While these mouse models provide approaches to study KSHV and EBV co-infection of B-cells in vivo, they do not recapitulate the complete infection pathways of these viruses, which utilize a lytic replication cycle in oropharyngeal epithelial cells.

The pathological effects of both KSHV and EBV are enhanced in immunosuppressed individuals, especially those co-infected with HIV exhibiting symptoms of AIDS [48, 49]. The impairment of the immune surveillance mechanisms is believed to reactivate latent infections by these oncogenic herpesviruses. KSHV and EBV are etiologically associated with AIDS-defining malignancies, including HIV-associated KS and non-Hodgkin lymphoma (NHL), where the KS tumor endothelial-like spindle cells and the NHL lymphoma cells are infected with KSHV and EBV, respectively. In addition, in AIDS-related PEL tumors, the majority of tumor cells are co-infected with KSHV and EBV [50]. We have shown that RFHV, the macaque RV1 homolog of KSHV is associated with simian AIDS-related retroperitoneal fibromatosis, a KS-like tumor that develops in associated with infections of simian immunodeficiency virus (SIV) or simian retrovirus 2 (SRV-2) [34, 51]. Furthermore, an increased incidence of lymphomas was observed in macaques experimentally infected with SIV [52] and the macaque LCV homologs of EBV have been detected in a large majority of SIV-associated lymphomas in different macaque species [53] [54–59]. Recently, we showed that simian AIDS-related lymphomas induced by either SIV or SRV-2 were etiologically associated with either MmuLCV or RRV in rhesus macaques, MneLCV or MneRV2 in pigtailed macaques and MfaLCV or MfaRV2 in cynomolgus macaques [60].

While experimental transmission of MmuLCV [61, 62] and RRV [63, 64] has been studied previously, no experimental transmission of RFHV, the macaque homolog of KSHV, has been reported. Despite more than a decade of efforts no lytic infection systems have been developed and no source of purified RFHV infectious particles has been identified. In the current study, we attempted to transmit RFHV using saliva from naturally infected macaques at the WaNPRC. We previously developed whole-virus multiplex luminex-based serological assays and qPCR assays for diagnosis of RV1 and RV2 rhadinovirus infections in macaques [65] [34, 51, 66]. We used these assays to screen the WaNPRC colony for seronegative/PCR-negative macaques as transmission recipients and RV1-positive macaques as transmission donors. Two negative juvenile macaques were identified as recipients and three adult macaques with high levels of salivary RV1 were identified as donors. QPCR and Luminex assays were used to follow experimental transmission over several years and various approaches were used to cause immunosuppression-induced viral reactivation. This preliminary transmission experiment resulted in the simultaneous co-infection of the macaque homologs of KSHV, EBV and HIV, with important similarities to transmission of the human viruses during the AIDS epidemic.

## Results

### Identification of RV1/RV2-naïve macaques

Initially, the macaque colony at the WaNPRC was screened for animals that were naïve to both macaque RV1 and RV2 rhadinoviruses to serve as recipients in a pilot experimental RFHV transmission. Sera from more than 200 macaques collected during the annual colony-wide health assessment screen were analyzed using a biplexed Luminex-based serological screen detecting macaque IgG antibodies reactive with KSHV virion proteins (RV1 assay, bead set 60) and RRV virion proteins (RV2 assay, bead set 61), as previously described [65]. The anti-RV1 antibody levels ranged from 9 to 2,425 MFI, with a median of 108 MFI, while the anti-RV2 antibody levels ranged from 4 to 2,705 MFI, with a median of 401 MFI (Table 1). Sixteen animals, of which 10 were identified as juveniles less than 13 months of age from the infant colony, had very low seroreactivities in both assays. The RV1 cutoff (mean of nonreactive juvenile sera plus 5 standard deviations) was determined to be 58 MFI, while the RV2 cutoff was 95 MFI [65]. High seroreactivity was observed in 45 macaques that were part of ongoing SIV vaccine studies with RV1 values ranging from 45 to 2,425 MFI with a median of 123 (40/45 positive samples) and RV2 values ranging 447–2,705 MFI with a median of 1,973 (44/45 positive samples) (Table 1)[65]. For comparison, the colony average (67 animals) was 108 MFI (RV1; range 49–448) and 350 MFI (RV2; range 163–1980), while the specific pathogen-free (SPF3; 37 animals) colony average was 99 MFI (RV1; range 30–465) and 475 MFI (RV2; range 42–1079) (Table 1). The SPF3 colony at the WaNPRC had been developed to eliminate cercopithecine herpesvirus 1 (Herpes B), SRV-2 and simian T-cell leukemia virus [67]. From the high RV1 and RV2 serological reactivities detected in the SPF3 colony, it was clear that the macaque RV1 and RV2 rhadinoviruses had not been eliminated from this colony. Two of the juvenile macaques, K04199 (0.5 year of age; RV1: 12 MFI, RV2: 6 MFI) and M04203 (1.0 year of age; RV1: 36 MFI, RV2: 42 MFI) were chosen as naïve recipients for the RFHV transmission experiment. PBMC were obtained from both animals and tested for RV1 and RV2 DNA using highly-sensitive and specific RV1 and RV2 qPCR assays [34, 66]. The assays were negative for both animals, further supporting their naïve status.

**Table 1.** Seroreactivity of Juvenile Pig-tailed Macaques

| Juvenile | Age (yr) | Median Fluorescent Intensity (MFI) |  |
| --- | --- | --- | --- |
|  |  | RV1 Virion (cutoff<sup>1</sup> = 58) | RV2 Virion (cutoff = 95) |
| K04027 | 0.5 | 11 | 5 |
| K04048 | 0.9 | 13 | 4 |
| K04183 | 0.7 | 84 | 45 |
| K04184 | 0.7 | 12 | 5 |
| K04199 | 0.5 | 12 | 6 |
| M04033 | 1.1 | 22 | 13 |
| M04038 | 1.0 | 11 | 5 |
| M04041 | 1.0 | 14 | 23 |
| M04205 | 0.5 | 9 | 8 |
| M04203 | 1.0 | 36 | 42 |
| **Positive Adult** |  |  |  |
| M03240 | 7.9 | 3674 | 1327 |
| M03126 | 8.3 | 2385 | 2239 |
| **Colony Average** |  |  |  |
| Colony<sup>2</sup> |  | 108 (49-448)<sup>3</sup> | 350 (163-1980)<sup>3</sup> |
| SPF3<sup>4</sup> |  | 99 (30-465)<sup>3</sup> | 475 (42-1079)<sup>3</sup> |
| Vaccine<sup>5</sup> |  | 123 (45-2425)<sup>3</sup> | 1973 (447-2705)<sup>3</sup> |

<sup>1</sup>Cutoff = mean of nonreactive juvenile sera plus 5 standard deviations
<sup>2</sup>Colony = general WaNPRC colony born and raised animals
<sup>3</sup>range of MFI
<sup>4</sup>SPF3 = animals from a specific pathogen-free colony at the WaNPRC that had been screened for the absence of simian retrovirus 2, simian T-cell leukemia virus and cercopithecine herpesvirus 1 (the macaque homolog of herpes simplex virus)
<sup>5</sup>Vaccine = animal in SIV vaccine studies

### Identification of RFHV-positive saliva donors

To identify RFHV-positive saliva donors, multiple saliva samples had been screened from 53 individual macaques at the WaNPRC over 3 years. The macaque population consisted of a mixture of pig-tailed macaques carrying RFHVMn and cynomolgus macaques carrying RFHVMf. Because high rhadinovirus serological reactivity was observed in macaques in ongoing SIV vaccine studies, saliva samples were obtained during regular testing in these studies. The saliva samples were tested for the presence of oral RV1 and RV2 DNA using specific RV1 and RV2 qPCR assays. While the majority of macaques never exhibited detectable levels of oral RV1 and RV2 rhadinovirus DNA in saliva, subsets were positive for oral RV1 (15/53) and RV2 (25/53). The maximal rhadinovirus levels ranged from 104–107 RV1 genomes/mL saliva and 103–108 RV2 genomes/mL saliva (Fig 1A). Fourteen of the 25 RV2 positive macaques had no evidence of oral RV1 DNA, while four of the 15 RV1 positive macaques showed no evidence of oral RV2 DNA (Fig 1B). Two cynomolgus macaques, 01019 and 98039. and one pig-tailed macaque, A02241, were identified as potential RFHV donors. During the course of the screening, all three of these macaques had high levels of RV1 DNA in saliva (Fig 2). Variable levels of oral RV2 DNA were occasionally detected, which were in most cases two logs or more less than RV1.

**Fig 1.**
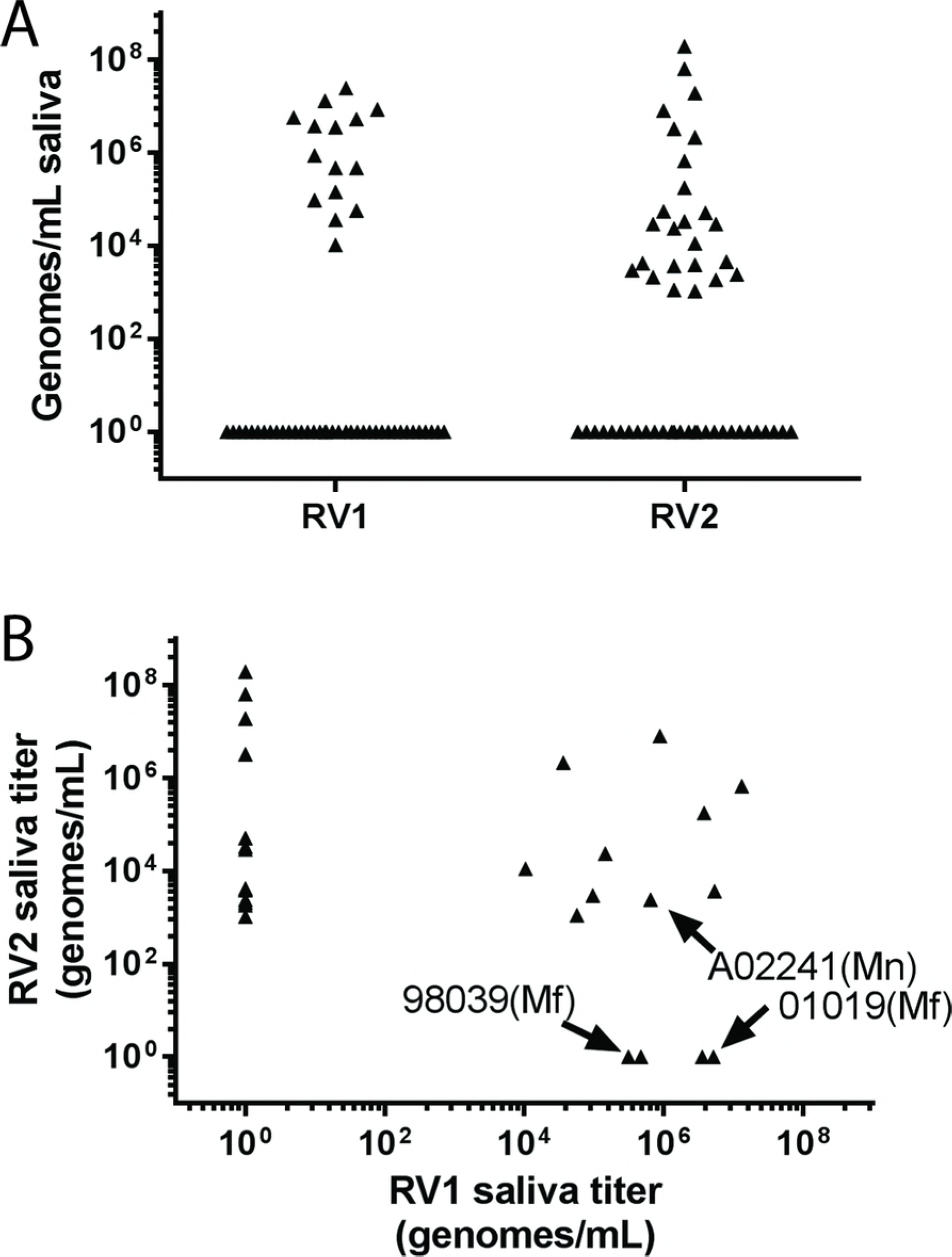
qPCR Screen for RV1 and RV2 Rhadinovirus DNA in Saliva. Saliva was obtained from macaques at the WaNPRC undergoing routine health screening and tested for the presence of RV1 and RV2 rhadinovirus DNA. A) The highest RV1 and RV2 DNA levels detected during longitudinal screening. B) Correlation of RV1 and RV2 viral loads for the animals in panel A showing a positive level of DNA for either RV1 or RV2.

**Fig 2.**
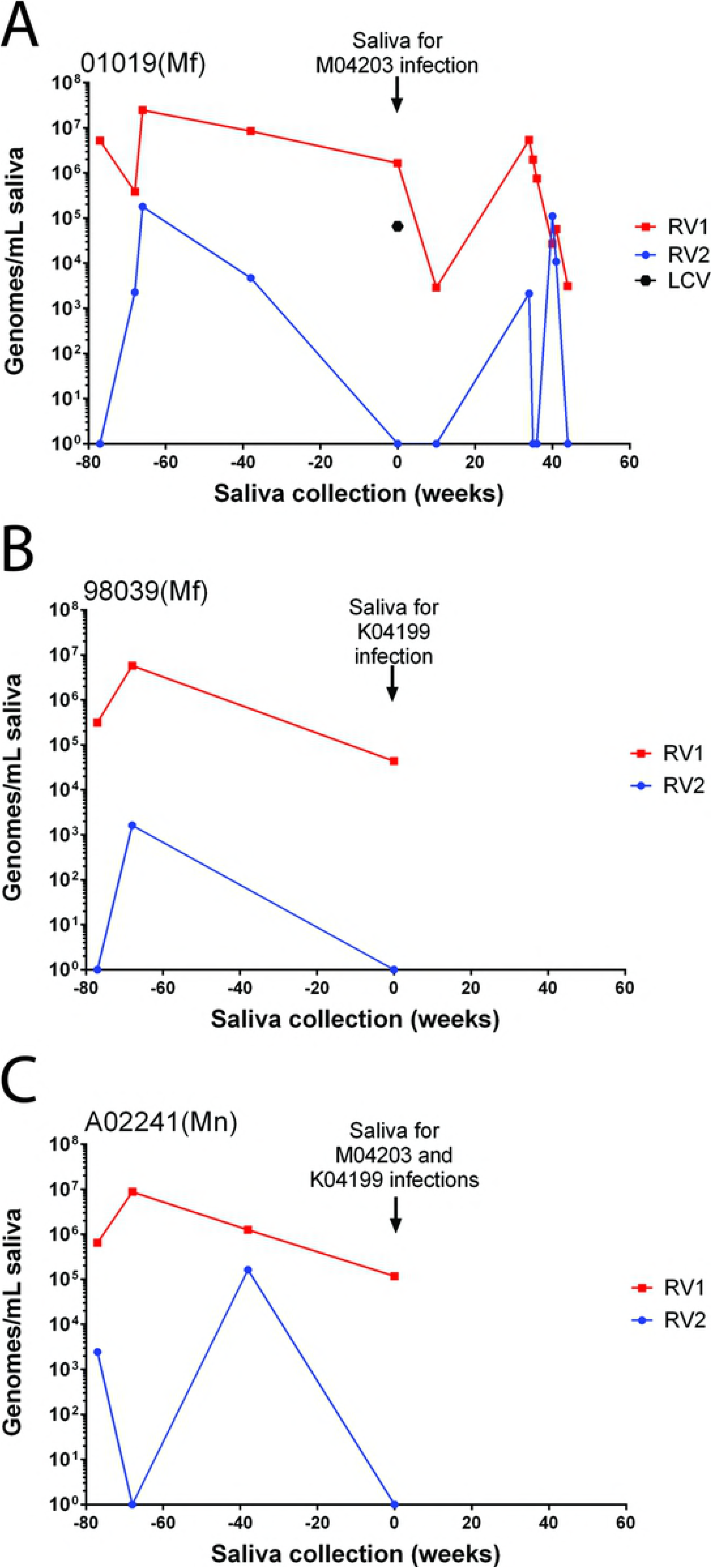
Longitudinal Analysis of RV1 and RV2 Rhadinovirus DNA in Saliva of Donor Macaques. The RV1 and RV2 DNA levels in longitudinal saliva samples of donor macaques A) cynomolgus macaque 01019, B) cynomolgus macaque 98039 and C) pig-tailed macaque A02241. Saliva collected on week 0 was used in the experimental transmission studies, as indicated. The level of LCV in the week 0 saliva of 01019 is shown in panel A.

### Experimental infection of naïve juvenile macaques with RFHV-positive saliva

Saliva was obtained from the two cynomolgus donors, 01019 (saliva “A”) and 98039 (saliva “C”). The 01019 and 98039 saliva samples contained 1.7 × 106 and 4.4 × 104 RV1 genomes per ml of saliva, respectively, with no detectable levels of RV2 DNA (Fig 2A, B). The 01019 saliva sample was also screened with a qPCR assay developed to detect macaque lymphocryptoviruses (LCV) [60], and was found to contain 6.6 × 10^4^ LCV genomes per ml (Fig 2A). The saliva was prepared as described in Materials and Methods and injected intravenously into the naïve juvenile pig-tailed recipients M04230 (01019 saliva “A”) and K04199 (98039 saliva “C”). Saliva was also obtained from the pig-tailed macaque donor A02441 (saliva “B”), which contained 1.2 × 10^5^ RV1 genomes per ml. This saliva was inoculated directly into the cheek pouches of both juvenile recipients (Fig 3). Thus, intravenously injected saliva would contain cynomolgus macaque viruses, while the cheek pouch inoculated saliva would contain pig-tailed macaque viruses. Saliva and blood samples were collected from the experimentally infected macaques weekly and/or bi-weekly and blood cells were counted. DNA was isolated from both saliva and peripheral blood mononuclear cells (PBMC) and RV1 and RV2 viral loads were determined by qPCR.

**Fig 3.**
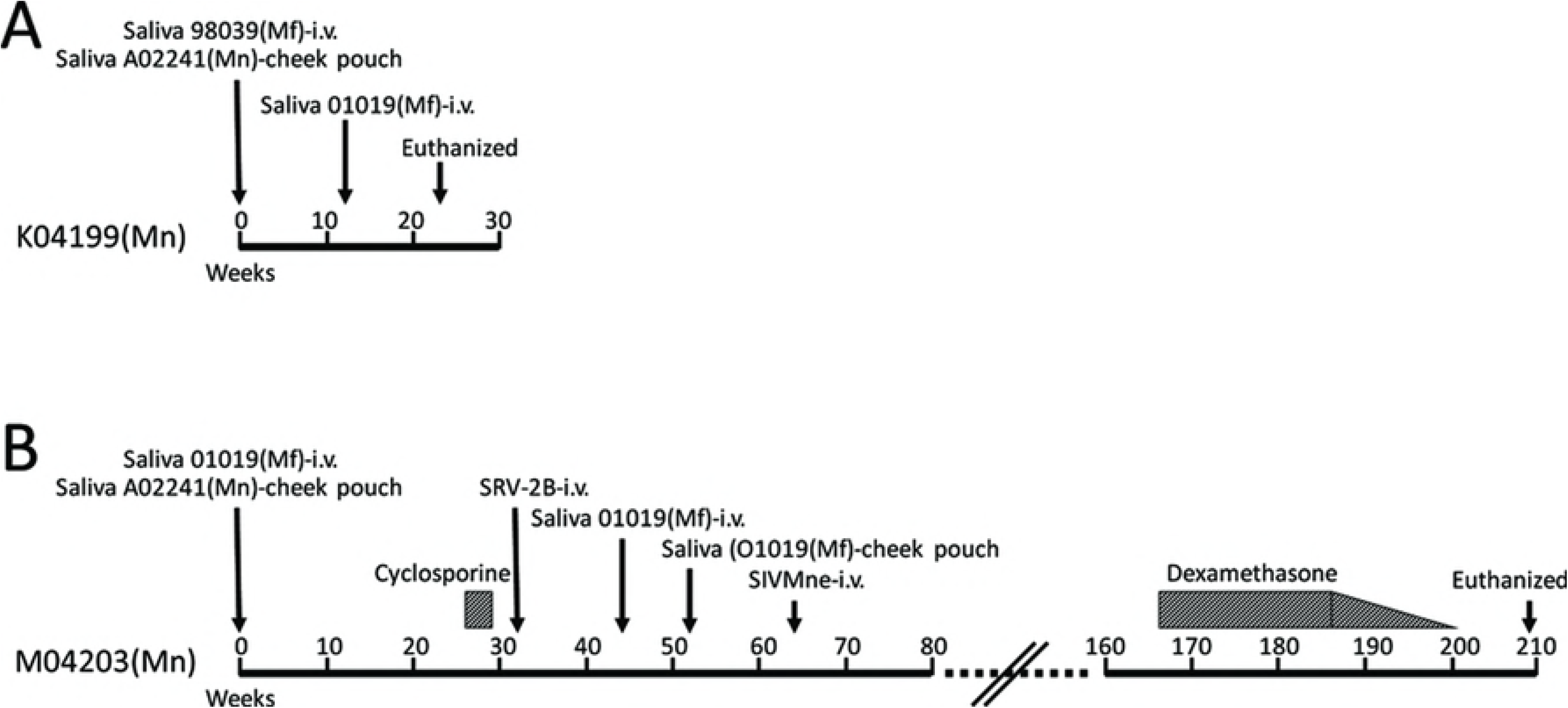
Timeline of Experimental Infections. The outlines of the experimental infections and treatment is shown for naïve juveniles A) K04199 and B) M04203, as described in the text and subsequent figure legends.

### Experimental transmission of salivary RFHVMn and SIV to K04199

The K04199 juvenile pig-tailed macaque was injected i.v. with cell-free saliva “C” from the cynomolgus macaque 98039 that had high titers of RFHVMf DNA and received whole saliva “B” into the cheek pouches from the pig-tailed macaque A02441 that had high titers of RFHVMn DNA. Analysis of saliva by RV1 qPCR revealed a low-level signal (153 RV1 genomes/ml saliva) by 10 weeks post infection (Fig 4A). The RV2 qPCR was negative during this time frame (Fig 4B). To enhance the infection, K04199 received an additional i.v. injection of cell-free saliva “A” from the cynomolgus macaque 01019, which had high titers of RFHVMf DNA in saliva. By week 16, the RV1 qPCR assay detected 2.3 million RV1 genomes/ml saliva. By week 18, however, the RV1 DNA levels were undetectable. The animal remained negative for RV2 during this time frame. Serological analysis of the bi-weekly blood samples revealed continuous baseline levels of anti-RV1 and anti-RV2 virion IgG antibodies from weeks 0 to 23 post infection (Fig 4A, B).

**Fig 4.**
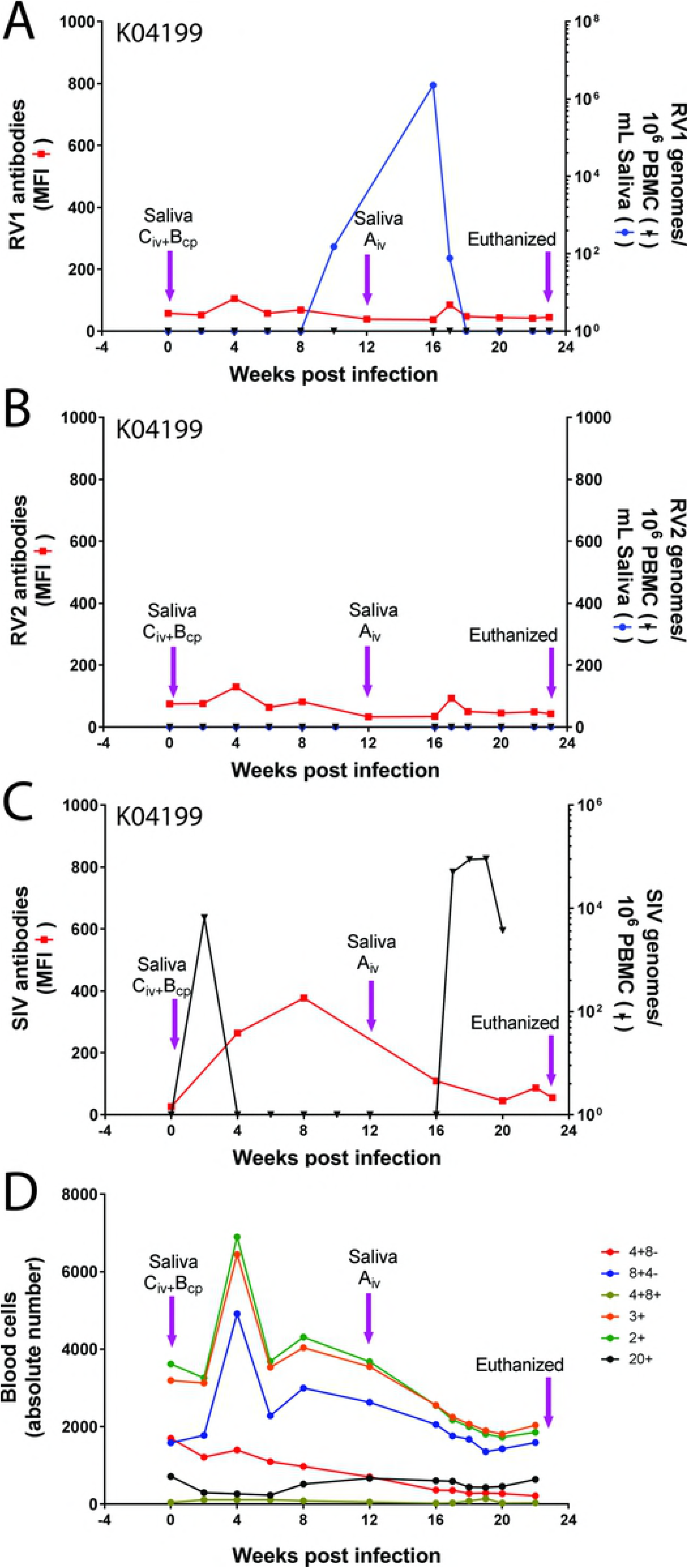
Experimental Rhadinovirus Infection in Juvenile Macaque K04199. The naïve juvenile pig-tailed macaque K04199 was inoculated intravenously with diluted, filtered saliva from the 98039 cynomolgus macaque donor (Saliva C) containing high levels of the RV1 rhadinovirus (RFHVMf), and was inoculated into the cheek pouches (cp) with whole saliva from the A02441 pig-tailed macaque (Saliva B) containing high levels of RFHVMn. Subsequently, K04199 was inoculated on week 12 with diluted, filtered saliva from the 01019 cynomolgus macaque donor (Saliva A) containing high levels of RFHVMf. Saliva and blood were collected weekly and/or biweekly. A) The RV1 genome levels were determined using the RV1 qPCR assay and the RV1 antibody levels were determined using the RV1 Luminex assay. B) the RV2 genome levels were determined using the RV2 qPCR assay and the RV2 antibody levels were determined using the RV2 Luminex assay. C) The SIV cDNA levels were determined using the SIV qPCR assay and the SIV antibody levels were determined using the SIV Luminex assay. D) The absolute number of cells in the lymphocyte subsets was determined, as indicated. K04199 developed clinical signs of SIV-induced AIDS and was terminated on week 23. MFI = mean fluorescent intensity.

The amplification product from the RV1 qPCR assay on the week 16 saliva sample was sequenced yielding a 100% match with the RFHVMn sequence and an 87% match with the RFHVMf sequence. These results indicate that the K04199 pig-tailed macaque had been experimentally infected with RFHVMn, the pig-tailed macaque RV1 rhadinovirus, from the A02441 whole saliva that had been inoculated into the cheek pouches of K04199. No evidence of an RFHVMf infection from the i.v. injection of cell-free saliva from the cynomolgus macaque donors 98039 or 01019 was detected.

Since the donor macaques were all long-term non-progressors in previous SIV vaccine studies and had been challenged with SIVmneB3718, we tested the longitudinal series of PBMC from K04199 for the presence of SIV using a SIV-specific qPCR assay [68]. A peak of SIV DNA (6,600 SIV genomes/10^6^ PBMC) was detected 2 weeks after the initial saliva infection. The SIV levels were undetectable by week 4 and remained undetectable until week 17, at which time the SIV DNA level increased to 51,000 SIV genomes/10^6^ PBMC. SIV DNA remained at this level for 2 weeks at which time K04199 showed symptoms of simian AIDS. Using a Luminex-based serological assay for SIV [67], low levels of IgG antibodies to SIV virion proteins were detected by week 4 (264 MFI), peaking at week 8 (377 MFI) and dropping to baseline levels thereafter. By week 23, K04199 developed clinical signs and hematological changes meeting the end-point criteria for AIDS protocols, and euthanasia was performed. Necropsy and histopathological examination revealed AIDS-associated pathologies, including involution of lymphoid tissues, notably thymus and spleen, pneumonia due to *Pneumocystis* sp. and orchitis. Blood counts showed a continual drop in the level of CD4^+^/CD8^−^ T-cells from 1,700 cells/ml on Day 0 to 212 cells/ml on Day 22 (Fig 4D).

### Experimental transmission of salivary RFHVMf to M04203

The M04203 juvenile pig-tailed macaque was injected i.v. with cell-free saliva “A” from the cynomolgus macaque 01019 that had high titers of RFHVMf DNA and received whole saliva “B” from the pig-tailed macaque A02441 that had high titers of RFHVMn DNA (Fig 3B). Analysis of M04203 saliva and PBMC by RV1 qPCR 4 weeks post infection revealed a low level of RV1 DNA in saliva (1,300 RV1 genomes/ml saliva) and a much greater level in PBMC (319,000 RV1 genomes/10^6^ cells) (Fig 5A). RV1 DNA in saliva and PBMC from weeks 6–10 post infection was below the level of detection. By week 16, RV1 DNA levels increased to 2,300,000 RV1 genomes/ml saliva with 27,000 RV1 genomes/10^6^ cells in PBMC. The high saliva levels maintained for two weeks, while the PBMC levels dropped to an undetectable level by week 17. During the first 16 weeks, the anti-RV1 virion antibodies remained at baseline levels but started rising sharply by week 18, reaching 3,300 MFI by week 20 (Fig 5A). During this timeframe and for the rest of the experiment, no RV2 genomes were detected by qPCR in either saliva or PBMC, and no specific anti-RV2 virion antibody response was observed. The amplification product from the RV1 qPCR assay on the week 16 saliva sample was sequenced yielding a 100% match with the RFHVMf sequence and an 87% match with the RFHVMn sequence. These results indicate that the M04203 pig-tailed macaque had been experimentally infected with RFHVMf following i.v. injection of the cynomolgus macaque RV1 rhadinovirus from the cell-free saliva of 01019. From weeks 20 to 26, the RV1 qPCR assays of both PBMC and saliva were negative and the anti-RV1 virion antibody level remained high, above 3,500 MFI.

**Fig 5.**
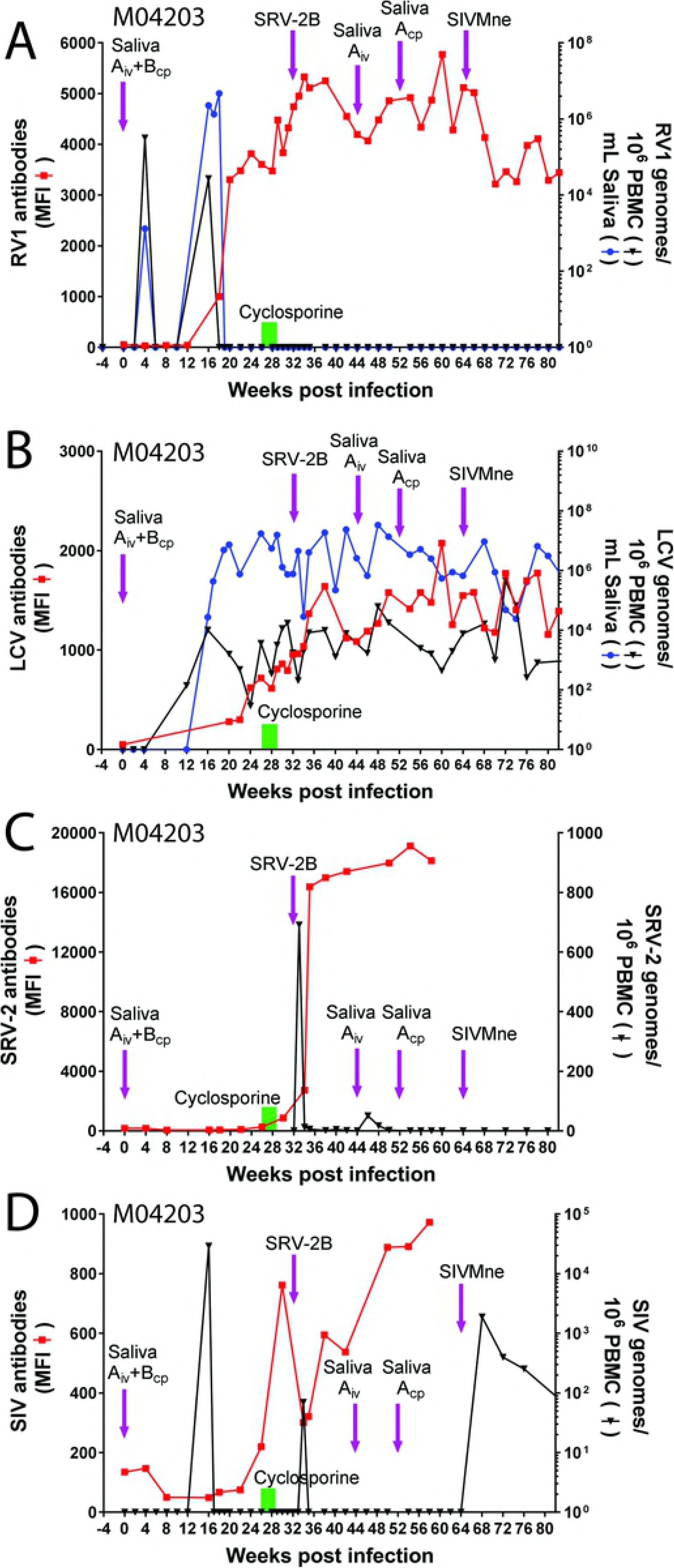
Experimental Rhadinovirus Infection in Juvenile Macaque M04203. The naïve juvenile pig-tailed macaque M04203 was inoculated intravenously (i.v.) with diluted filtered saliva from the 01019 cynomolgus macaque donor (Saliva “A”) containing high levels of RFHVMf and low levels of MfaLCV and was inoculated into the cheek pouches with whole saliva from pig-tailed macaque A02241 (Saliva “B”), which had high levels of RFHVMn. M04203 received a second iv inoculation of saliva from 01019 (Saliva “A”) 44 weeks after the initial infection. Subsequently, whole saliva from 01019 (Saliva “A”) was administered to the cheek pouch at week 52. To activate rhadinovirus infections, M04203 was treated with daily injections of cyclosporine for 3 weeks (week 26–29) to induce immune suppression, as indicated (green bar). The animal was subsequently infected with SRV-2B and SIVMne, which have previously been associated with rhadinovirus activation and associated pathologies (see text). Saliva and blood were collected weekly and/or bi-weekly. A) The RV1 genome levels were determined using the RV1 qPCR assay and the RV1 antibody levels were determined using the RV1 Luminex assay. B) the LCV genome levels were determined using the LCV qPCR assay and the LCV antibody levels were determined using the LCV Luminex assay. C) The SRV2 cDNA levels were determined using the SRV2 qPCR assay and the SRV2 antibody levels were determined using the SRV2 Luminex assay. D) The SIV cDNA levels were determined using the SIV qPCR assay and the SIV antibody levels were determined using the SIV Luminex assay.

### Identification of MfaLCV co-infection in M04203

To determine whether M04203 had been co-infected with LCV from the 01019 saliva, the saliva and PBMC samples from M04203 were screened with the LCV qPCR assay. LCV DNA was detected in the PBMC 12 weeks after the initial saliva infection (142 LCV genomes/10^6^ PBMC) and in the saliva 16 weeks after infection (27,438 LCV genomes/ml saliva) (Fig 5B). LCV DNA was continually detected in PBMC and saliva reaching 3.6 × 10^3^ LCV genomes/10^6^ PBMC and 1.7 × 10^7^ genomes/ml saliva by week 26. The amplification product from the LCV qPCR assay on the week 16 saliva sample was sequenced yielding a 100% match with the MfaLCV sequence. These results indicate that the M04203 pig-tailed macaque had been experimentally co-infected with both RFHVMf, the cynomolgus macaque RV1 rhadinovirus, and MfaLCV, the cynomolgus macaque LCV, via the i.v. injected cell-free saliva “A” sample from 01019. To further study the LCV infection, we developed a Luminex-based serological assay to detect macaque antibodies that cross-react with EBV virion proteins, as described in Materials and Methods. No anti-LCV virion antibodies were detected at day 0, but by week 26, low levels of anti-LCV antibodies were observed (720 MFI) (Fig 5B).

### Identification of SIV co-infection in M04203

Since SIV was detected in K04199 from the saliva inoculation, which resulted in high levels of SIV DNA by week 17 and clinical signs of AIDS by week 23, we screened the PBMC samples of M04203 for evidence of a saliva-induced SIV infection. High levels of SIV DNA (2.9 × 10^4^ genomes/10^6^ PBMC) were detected in PBMC at week 16, coinciding closely with the appearance of SIV DNA in K04199 at week 17. However, unlike K04199, which continued to have high levels of SIV DNA in PBMC from week 17 to week 23, at which time it was sacrificed with clinically evident AIDS, M04203 had no detectable SIV DNA during this timeframe (Fig 5A) and showed no evidence of AIDS-associated clinical signs. Analysis of blood cells revealed consistent levels of circulating B-cells (CD20+) and T-cells (CD4+8−, CD8+4−, CD4+8+, CD3+, CD2+) through week 16 (Fig 6). However, immediately after the appearance of RFHVMf and MfaLCV DNA in saliva and PBMC and SIV DNA in PBMC on week 16, the absolute number of B-cells and T-cells dropped precipitously reaching a minimum on week 20 (Fig 6), at which time the strong anti-RV1 virion antibody and weaker anti-LCV antibody responses initiated (Fig 5A).

**Fig 6.**
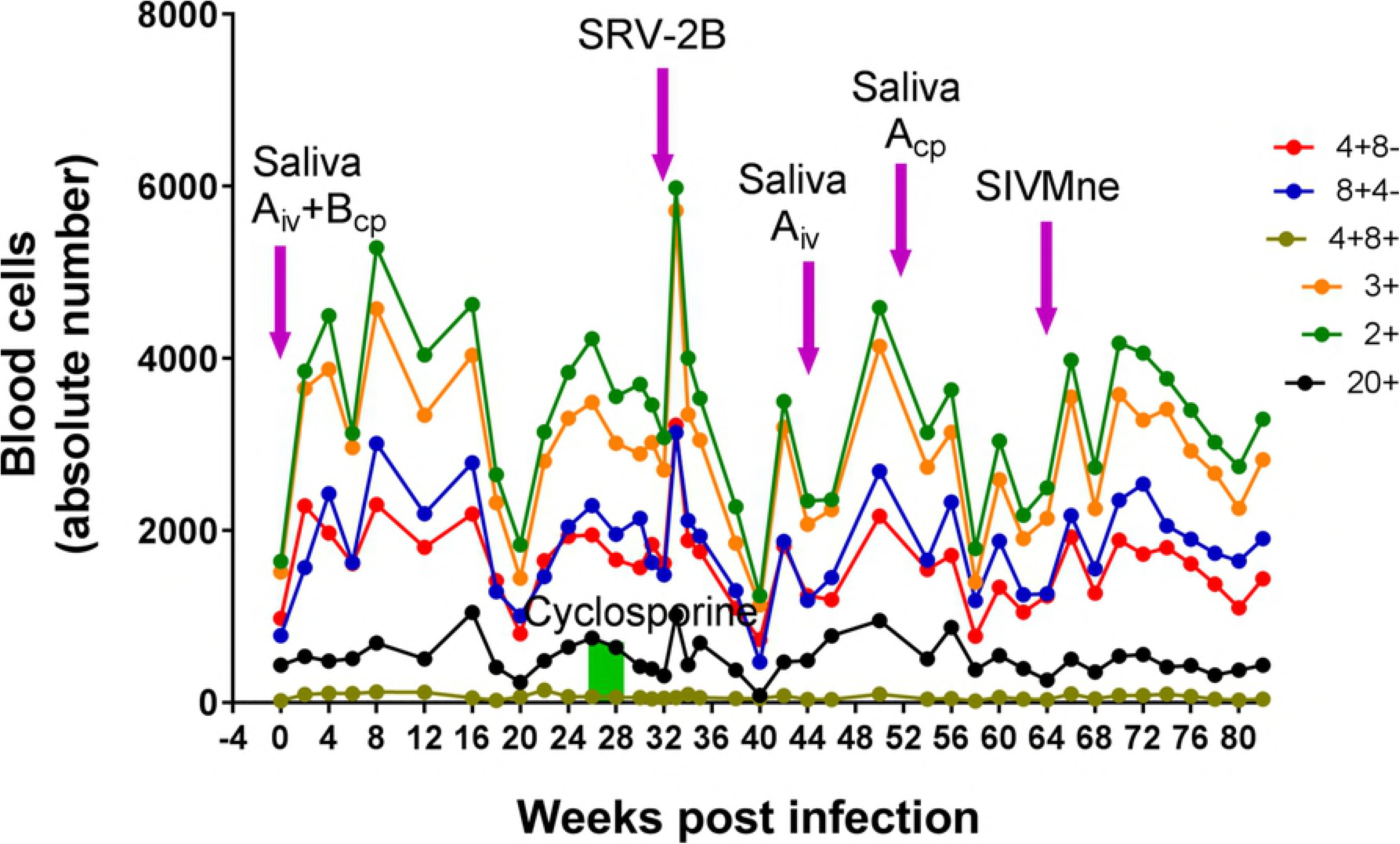
Longitudinal Analysis of Lymphocyte Subsets in M04203. The absolute number of cells in lymphocyte subsets was determined from the blood samples in the experimental infection of M04103 shown in Fig 5.

### Cyclosporine-induced immunosuppression of M04203

Previously, cyclosporine has been used to immunosuppress macaques, resulting in a high incidence of lymphomas and post-transplant lymphoproliferative disorders associated with macaque homologs of EBV [54, 69–71]. To examine the effect of immunosuppression on the RFHVMf and MfaLCV infections, M04203 was treated daily from weeks 26–29 with cyclosporine A intramuscularly at 15 Mgs/Kg (Fig 3B). Although no RV1 DNA was detected in PBMC or saliva from week 26–32, the titer of anti-RV1 virion antibodies continued to rise increasing from 3,607 (week 26) to 4,741 MFI (week 32) (Fig 5A). This suggests that the immunosuppressive effect of cyclosporine on T-cells may have caused a low-level reactivation of the latent RFHVMf infection resulting in the expression of sufficient RFHMf virion proteins to increase the level of anti-virion antibodies, without evidence of DNA replication. While the level of antibodies to MfaLCV virion proteins continued to increase during this timeframe, no obvious effects were observed on the levels of LCV DNA in saliva or PBMC, which continued to be high (Fig 5B). A sharp peak of anti-SIV antibodies was detected after the cyclosporine treatment reaching a maximum at week 30 (762 MFI) (Fig 5D).

### Co-infection of M04203 with SRV-2B

Previous studies have shown that simian retrovirus 2 (SRV-2) infection has an immunosuppressive effect on macaques and a high percentage of naturally infected animals at the WaNPRC developed high levels of RV1 and RV2 rhadinovirus infections in conjunction with rhadinovirus-associated malignancies [34, 60, 72, 73]. To examine the effect of SRV-2 on the RFHVMf and MfaLCV infections, M04203 was injected i.v. on week 32 with infectious SRV-2B (1 × 10^7^ viral genomes) that had been isolated from a naturally infected macaque (T81273) at the WaNPRC (Fig 3B). The SRV-2B strain of SRV-2 was previously associated with the long-term development of RFHV-associated retroperitoneal fibromatosis (RF) [74]. An SRV-2B-specific qPCR assay was developed to quantitate viral loads (see Materials and Methods), and antibodies to SRV-2 were quantitated using a Luminex-based assay developed previously at the WaNPRC [67]. One week after the SRV-2B infection (week 33), SRV-2B cDNA was detected in PBMC (691 SRV-2B genomes/10^6^ PBMC) (Fig 5C). Anti-SRV-2 IgG antibodies (2,737 MFI) were detected one week later (week 34) and increased to 16,386 MFI by week 36. Anti-SRV-2 antibody levels continued to increase reaching a maximum of 19,118 MFI at week 54. With the exception of minimal signals at weeks 46 and 48, SRV-2B cDNA was below the level of detection in PBMC (Fig 5C). The RV1 and RV2 qPCR assays remained negative during this time frame (Fig 5A). The anti-RV1 antibody levels continued to increase from week 28 reaching a maximum of 5,331 MFI by week 34 and dropped to 4,194 MFI by week 44 (Fig 5A). The LCV DNA levels in saliva and PBMC remained consistently high during this timeframe, while the anti-LCV antibody levels reached a maximum of 1,642 MFI at week 38 and decreased to 1,122 MFI by week 44. Two weeks after the SRV-2B infection (week 34), the anti-SIV antibody levels had dropped to 300 MFI and a low level of SIV DNA was detected (70 SIV genomes/10^6^ PBMC). Subsequently, the anti-SIV antibody levels increased to 973 MFI (week 58) and SIV DNA was no longer detected.

### Secondary saliva infections of M04203

In attempts to further augment the herpesvirus infections in M04203, additional saliva “A” from 01019 was inoculated i.v. at week 44 and into the cheek pouch at week 52 (Fig 3B). While the anti-RV1 virion antibody level remained high after these treatments, reaching a maximum of 5,771 MFI (week 60), no RV1 or RV2 DNA was detected in PBMC or saliva (Fig 5A). The LCV DNA and antibody levels remained consistently high. The anti-SIV antibody levels continued to rise and no SIV DNA was detected in PBMC.

### Co-infection of M04203 with SIVMne

Previous studies have shown that SIV infection has potent immunosuppressive effects on macaques resulting in the development of simian AIDS-associated malignancies, which is characterized by high levels of rhadinovirus and/or lymphocryptovirus in the tumor lesions [51, 60]. Since SIV DNA was no longer detected in PBMC at week 62, M04203 was injected i.v. with high-titer purified SIVMne on week 64 to try to induce a lentivirus-associated immunosuppression. High levels of SIV DNA were detected after the SIV inoculation reaching a maximum (1,887 SIV genomes/10^6^ PBMC) four weeks after inoculation (week 68) (Fig 5D). The level of SIV declined to 63 SIV genomes/10^6^ PBMC by week 82. The RFHVMf DNA levels in saliva and PBMC remained undetectable after the SIVMne infection and no obvious changes in LCV status were noted, as LCV DNA and anti-LCV antibody levels remained high.

### Quantitation of blood lymphocytes in M04203

The M04203 blood lymphocytes were immunophenotyped and quantitated in the longitudinal samples. An increase in the various T-cell subsets was observed 2 weeks after the initial saliva inoculation (Fig 6A). Four weeks after the appearance of high levels of RFHVMf, MfaLCV and SIV at week 16, a dramatic reduction in the level of the T-cell and B-cell subsets was observed, which rebounded by week 26. A second increase in the circulating T-cell subsets was detected 2 weeks after the experimental SRV-2B infection. This increase was followed by a dramatic reduction in the T-cell and B-cell subsets 7 weeks after the SRV-2B infection (week 40). The T-cell and B-cell populations again rebounded quickly showing some variation through week 82 (Fig 6A), which did not show any correlation with saliva inoculations or SIVMne infection. Throughout the experimental time frame, the level of CD4^+^CD8^−^ T-cells remained high (Fig 6), showing no evidence of lentivirus induced immunosuppression, contrasting to the situation observed in K04199 (Fig 4D).

### Long-term monitoring of M04203 and dexamethasone treatment

Blood and saliva samples of M04203 were continually collected for 209 weeks (~ 4 years) after the initial saliva inoculation, at which time the experiment was terminated and the animal was culled. Prior to termination, M04203 was treated with dexamethasone 2 mg/kg/day for 7 days (weeks 164–165) and then with 1 mg/kg/day every three days (week 165 to week 179) after which the treatment tapered 10% per week until the dose was 0.1 mg/kg/day by week 201 (see Fig 3). Dexamethasone is a glucocorticoid agonist that has multiple anti-inflammatory and immunosuppressive functions [75]. The CD4/CD8 ratio of M04203 remained relatively constant (~ 0.8) until the initiation of dexamethasone treatment at week 164, at which point the ratio dropped precipitously reaching a level of 0.08 by week 176 (Fig 7D). The CD4/CD8 ratio varied between 0.1 and 0.2 during the treatment period and rose slightly by week 209 (0.35). Until week 164, the total neutrophil and monocyte counts remained constant while the lymphocyte counts decreased slowly to 50% of the initial values (Fig 7C). Dexamethasone treatment induced high levels of circulating neutrophils, which returned to baseline after treatment, while a concomitant and significant drop in lymphocyte counts was observed (Fig 7C). The lymphocyte numbers rebounded during the later stage of the tapered drug treatment and the monocytes showed a transient increase.

**Fig 7.**
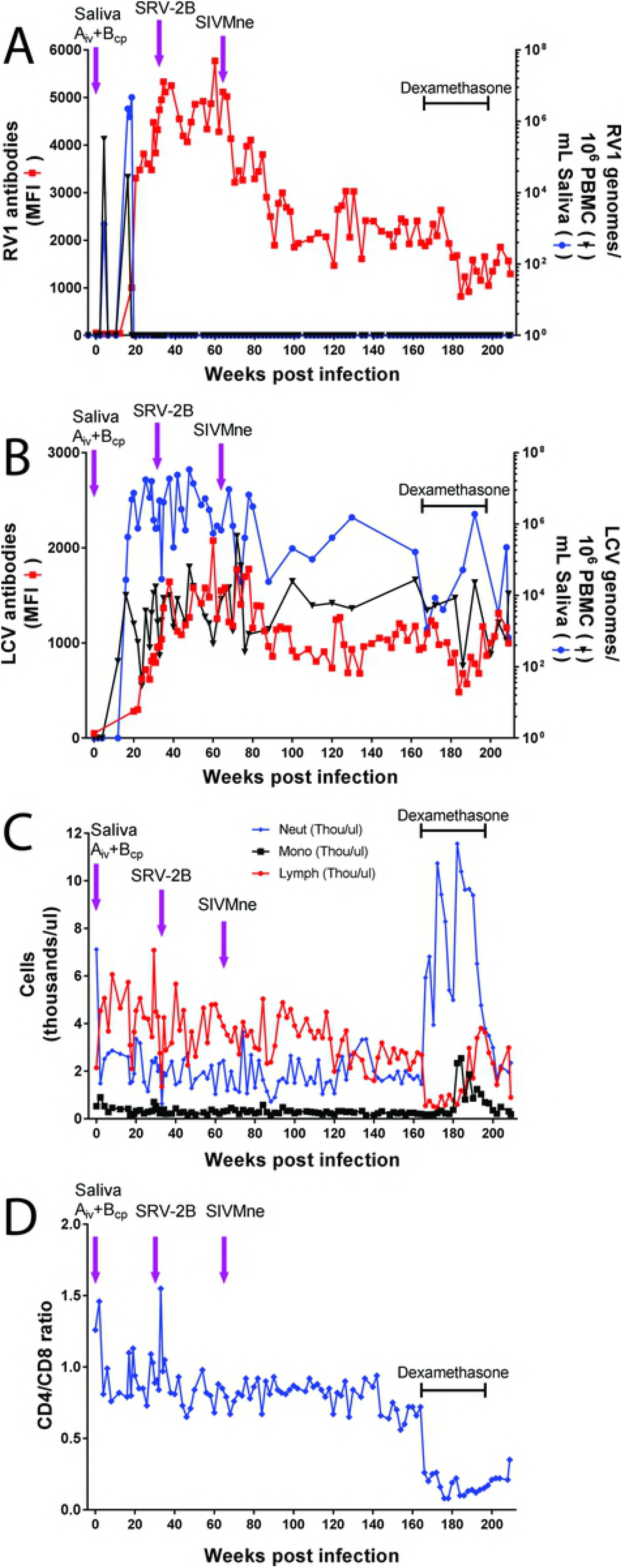
Complete Longitudinal Analysis of the Experimental Infection of M04203. The juvenile macaque M04203 was experimentally inoculated with saliva from the donor macaques as indicated in the legend to Figure 5. Blood and saliva samples were collected weekly and/or biweekly until the experiment was terminated 209 weeks after the initial saliva inoculation and M04203 was culled. At week 164, the animal received a course of dexamethasone treatment as indicated in the text and Figure 3. A) The RV1 genome levels were determined using the RV1 qPCR assay and the RV1 antibody levels were determined using the RV1 Luminex assay. B) the LCV genome levels were determined using the LCV qPCR assay and the LCV antibody levels were determined using the LCV Luminex assay. C) The absolute count of neutrophils, monocytes and lymphocytes was determined. D) the CD4/CD8 ratio was determined from the lymphocyte count.

The effects of dexamethasone on the blood cell populations had no obvious effect on the RFHVMf infection, as no RV1 DNA was detected in saliva or PBMC during the remainder of the experiment (Fig 7A). The high levels of anti-RV1 virion antibodies detected during the first 80 weeks after the initial saliva inoculation slowly decreased achieving a relatively stable level of ~2,000 MFI during the next 100 weeks (Fig 7A). The RV1 antibody levels dropped to 800 MFI during the dexamethasone treatment period and restabilized at ~1,500 MFI after treatment. From weeks 80 to 180, the levels of LCV in saliva and PBMC and the levels of anti-LCV virion antibodies remained high (Fig 7B). Dexamethasone treatment decreased the anti-LCV antibody levels, which rebounded similar to that seen with the anti-RV1 antibodies. An initial drop in the level of LCV DNA in saliva was observed after the dexamethasone treatment (1.7 × 10^5^ to 1.1 × 10^3^ LCV genomes/ml saliva). The LCV DNA level rebounded to 2 x 10^6^ LCV genomes/ml saliva during the tapering stages of the dexamethasone treatment but dropped to 600 at week 209. At this time, the level of LCV DNA in PBMC was equivalent to the level before dexamethasone treatment (~10^4^ LCV genomes/10^6^ PBMC).

At necropsy the tissues and organs of M04203 were grossly unremarkable, with no evidence of disease processes. Analysis by qPCR revealed no evidence of RV1 or RV2 DNA in saliva, PBMC, ileocecal junction, liver, mesenteric lymph nodes, salivary gland, spleen or tonsil (Table 2). In contrast, LCV DNA was detected in both saliva (657 LCV genomes/ml saliva) and PBMC (11,049 LCV genomes/10^6^ PBMC) at the time of necropsy and low levels of LCV were detected in the tonsil, mesenteric lymph node, salivary gland, and spleen (Table 2).

**Table 2.** Viral load in M04203 Necropsy Tissue

| Tissue | Viral genomes per $10^6$ cells | | |
| --- | --- | --- | --- |
|  | RV1 | RV2 | LCV |
| PBMC | Nd<sup>1</sup> | Nd | 11,049 |
| Lymph node (mesenteric) | Nd | Nd | 27 |
| Spleen | Nd | Nd | 1 |
| Salivary gland | Nd | Nd | 10 |
| Tonsil | Nd | Nd | 1,215 |
| Ileocecal junction | Nd | Nd | Nd |
| Liver | Nd | Nd | Nd |

<sup>1</sup> Nd = not detected

## DISCUSSION

Although RFHV infection in macaques represents a close animal model of KSHV biology and pathology, the development of an experimental model using RFHV has lagged due to the lack of a source of infectious virus. Our pilot study was designed to determine whether experimental transmission of RFHV could be achieved using saliva inoculations from RFHV-infected donors into rhadinovirus-naïve recipients. Previous studies showed that the majority of macaques in captive populations are co-infected with rhadinoviruses belonging to both the RV1 and RV2 lineages [65, 76]. While specific pathogen-free colonies lacking both of these viruses have not been developed at the WaNPRC or other national primate centers, we identified two hand-reared juvenile pig-tailed macaques that were naïve to both RV1 and RV2 rhadinoviruses by serological and molecular methods. We previously identified a cohort of possible donor macaques from ongoing SIV/SHIV vaccine studies at the WaNPRC that had high titers of anti-RV1 antibodies [65]. Laborious screening of these animals identified two cynomolgus and one pig-tailed macaque as possible donors that had high levels of RV1 and low levels of RV2 in saliva. While transmission of a pig-tailed macaque RV1 rhadinovirus (RFHVMn) to a naïve pig-tailed macaque recipient would represent a homologous infection system, the two cynomolgus macaques carrying the cynomolgus RV1 rhadinovirus (RFHVMf) had higher levels of RV1 rhadinovirus and lower levels of RV2 rhadinovirus providing an increased chance for a singular cross-species RV1 transmission. We therefore designed a protocol to inoculate saliva containing RFHVMf from each of the two cynomolgus macaques into the naïve pig-tailed macaques by iv injection and to inoculate whole saliva containing RFHVMn from the single pigtailed macaque into the cheek pouches of both naïve pig-tailed macaques. Since RFHVMn and RFHVMf could be distinguished by sequence analysis, this provided the best option to confirm RV1 transmission and determine the route of infection.

After the saliva inoculations, the RV1 qPCR assay was transiently positive in both juvenile recipients at week 16 post inoculation. Sequence analysis of the PCR products revealed the presence of the RFHVMn species in the K04199 pig-tailed macaque recipient indicating that the transmitting rhadinovirus was from the pig-tailed macaque donor A02441 through a natural oral route of infection with saliva. Although K04199 could already have been naturally infected with RFHVMn at the time of the saliva inoculation, there was no evidence of a prior infection either by PCR or serology. In contrast, the RV1 qPCR assay identified an RFHVMf infection in the M04203 pig-tailed juvenile recipient, indicating that the transmitting species was RFHVMf from the cynomolgus macaque donor 01019 through an i.v. exposure. Although the complete sequence of the cynomolgus RFHVMf has not been determined, strong sequence similarity has been detected between individual genes of the pig-tailed and cynomolgus RFHV species [22, 23], suggesting that cross-species RFHV infections could have similar biological and pathological outcomes. In addition, no differences in pathogenicity were observed in cross-species infection of the closely related rhesus and pig-tailed RV2 rhadinoviruses, RRV and MneRV2(PRV) [63].

Longitudinal analysis of blood and saliva samples from the inoculated saliva recipients showed no evidence of experimental transmission of an RV2 rhadinovirus to either juvenile macaque recipient. Although our experimental protocol was designed to induce a single experimental RFHV infection in the naïve macaques using high titer RFHV-positive macaque saliva, we detected LCV DNA in the saliva of our most optimal donor 01019. The screening of saliva, serum and PBMC from the infected M04203 recipient clearly showed evidence of a robust infection with MfaLCV from the 01019 cynomolgus macaque donor. Thus, both RFHVMf and MfaLCV had been co-transmitted i.v. from the saliva of the 01019 cynomolgus macaque donor to the M04203 pig-tailed macaque recipient. The appearance of RFHVMn in the saliva of K04199 matched the time frame for the appearance of RFHVMf and MfaLCV in the M04203 recipient, approximately 17 weeks post inoculation, suggesting that all three infections were experimental and that the time frame for establishing detectable infections through cheek pouch or i.v. inoculations were similar.

Since the RFHV-positive macaque donors had all been challenged previously with SIVmneB3718 in other, unrelated studies, the possibility existed that saliva inoculation of the juvenile recipients could transmit SIV. Since SIV infections have been shown to exacerbate gammaherpesvirus infections, these macaques represented an optimal source of saliva to demonstrate RFHV transmission. Our data showed that both M04203 and K04199 juveniles were infected with SIV from one of the donor saliva samples. High levels of SIV DNA replication intermediates were detected in PBMC of both animals by week 17 post inoculation. While a weak immune response to SIV was detected by week 4 in K04199, this decreased to a minimal level by week 17 at which time high levels of SIV DNA were continuously detected in PBMC and the animal developed clinical signs of simian AIDS. In contrast, the level of SIV DNA in M04203 dropped immediately after the peak at week 16 and the levels of anti-SIV antibody continued to increase during the initial year following saliva inoculation and no clinical signs of simian AIDS were observed. While RFHV infection was observed in both juvenile macaques, neither the uncontrolled SIV infection in K04199 nor the controlled SIV infection in M04203 resulted in any detectable activation or lytic replication of RFHV.

While both juvenile recipients showed evidence of experimental RFHV transmission by week 16 with RFHV DNA detected in saliva and/or PBMC, M04203 showed a strong IgG response to RFHV virion proteins, whereas no RFHV immune response was detected in K04199. K04199 showed a loss of T-cells, especially CD4+ cells, which coincided with the strong increase in SIV cDNA in PBMC, a low level of anti-SIV antibodies and development of simian AIDS. Thus, the K04199 recipient was unable to mount a strong immune response to RFHVMn, the natural RV1 rhadinovirus of pig-tailed macaques, presumably due to perturbations of the immune system by SIV. On the other hand, M04203 showed no loss of CD4+ T-cells throughout the experimental timeframe, even in the presence of ongoing SIV and SRV-2B infections, and cyclosporine or dexamethasone immunosuppressive treatments. In attempts to induce different types of immunosuppression to reactivate the RFHVMf infection, M04203 was sequentially treated with immunosuppressive agents, including cyclosporine, SRV-2 D-type retrovirus, SIVMne lentivirus and dexamethasone. No RFHVMf DNA was detected in saliva or PBMC after cyclosporine treatment, that would have indicated a viral reactivation. The antibody response to RFHV virion proteins continued to rise sharply after the cyclosporine treatment, suggesting continual immune stimulation by proteins associated with RFHVMf replication. The SRV-2B infection was followed by a peak of SRV2 cDNA in PBMC two weeks after the experimental SRV2 infection and a robust anti-SRV2 immune response that immediately controlled the SRV2 infection. The baseline CD4^+^/CD8^+^ ratio did not change after SRV-2 infection and no activation of the latent RFHV infection was observed. The subsequent experimental SIVMne infection was followed by a high level of SIV cDNA in PBMC. The baseline CD4^+^/CD8^+^ ratio did not change and no activation of the latent RFHVMf was observed. As a final step, dexamethasone-induced immunosuppression was used to try to induce RFHV reactivation. A large decrease in the CD4^+^/CD8^+^ lymphocyte ratio was detected with a concomitant increase in neutrophils. While this treatment decreased the anti-RV1 virion antibody level by 50%, no RFHVMf DNA was detected in saliva or PBMC indicating no activation of the rhadinovirus. During this treatment, the levels of MfaLCV were variable but remained high in both PBMC and saliva, indicating that the immunosuppressive treatments had little effect on the activated state of the lymphocryptovirus.

Recent studies in Uganda have investigated the role of co-infection on the shedding of KSHV and EBV in saliva in HIV-negative mothers and their children [77]. In this population, individuals were more likely to shed EBV than KSHV in saliva. An inverse relationship between the level of EBV and KSHV shedding was detected, suggesting a direct or indirect interaction between the two viruses. Other studies have shown that persistent KSHV infection increases EBV-associated tumor formation and enhances EBV lytic gene expression [47]. The infection status and outcome of M04203 co-infected with the macaque homologs of EBV and KSHV reflects the findings in the Uganda study by Newton et al. as high level lymphocryptovirus shedding was detected in the absence of rhadinovirus shedding. Even though the putative infective dose of MfaLCV was 250-fold lower than RFHVMf, MfaLCV established a persistent lytic infection with oral shedding, while RFHVMf showed little evidence of viral reactivation or replication. While this could be due to an effect of the persistent rhadinovirus infection on lymphocryptovirus lytic gene expression, as suggested previously [47], it could also be due to differences in the biology and tropism of the two viral lineages and differential control of lytic activation and replication by the immune system of the infected host.

Temporally, the cell-free salivary MfaLCV, RFHVMf and SIV infections in M04203 initiated simultaneously with nearly identical kinetics. Viral DNA was detected in PBMC in all three cases at the same time (~16 weeks post inoculation) indicating transmission to a population of circulating lymphocytes. Furthermore, both MfaLCV and RFHVMf DNA were detected in saliva indicating viral transmission and replication in oral epithelial cells and shedding of infectious virions in the same timeframe. The initial transmission and replication of RFHVMf and MfaLCV in M04203 occurred in the absence of detectable IgG anti-virion antibody responses. Subsequently, M04023 mounted strong long-term immune responses to the virion proteins of both RFHVMf and MfaLCV, indicating successful cross-species infection with the cynomolgus homologs of RFHV and LCV. Unlike many other serological assays for KSHV and EBV, the Luminex assays used in this study detect antibodies to virion proteins, which are generated during primary infections and reactivation of lytic replication and do not target antibodies to specific latency antigens [65]. No RFHV DNA was detected in PBMC or saliva of M04203 after week 19, indicating that the immune system was controlling the RFHVMf infection and suppressing viral reactivation. In contrast, high levels of MfaLCV DNA were detected in both saliva and PBMC of M04203 throughout the experiment, in spite of a robust anti-LCV immune response. Thus, simultaneous rhadinovirus and lymphocryptovirus infections resulted in very distinct outcomes under the same level of immune competency, with the rhadinovirus RFHVMf displaying a very strong latency phenotype and the lymphocryptovirus MfaLCV displaying an ongoing lytic phenotype. Clearly, more research on these aspects are warranted, but may have to await the availability of pure isolates of all these viruses to allow controlled homologous co-infections.

## Materials and methods

### Cells

Raji cells (P8) were obtained from M. Thouless (UW) and cultured in RPMI at 37 ^o^C.

### Animals

All macaques (*M. nemestrina* and *M. fascicularis*) were housed at the Washington National Primate Research Center (WaNPRC) from 2005–2009 in accordance with standards of the American Association for Accreditation of Laboratory Animal Care and the experimental protocol3146-03 was approved by the University of Washington Institutional Animal Care and Use Committee. The animals were restricted to protected contact due to veterinary and research staff concerns regarding potential injury and spread of infection. They were fed a nutritionally balanced diet of monkey biscuits twice per day, participated in the WaNPRC Environmental Enhancement Plan, and were provided enrichment items on a daily basis. The research adhered to the American Society of Primatologists Principles for the Ethical Treatment of Nonhuman Primates. All animals were housed indoors on a 12-hr light cycle. The subjects were housed in two-tiered stainless steel cages, complying with Animal Welfare Act USDA standards for NHPs based on animal weight. The animals were observed daily by trained animal care staff, and underwent ketamine-sedated physical examination including CBC/serum chemistry monthly.

Blood and saliva samples were obtained under anesthesia using Ketamine. Experimental endpoints were in accordance with the University of Washington IACUC-approved AIDS Monitoring Protocol, which is a scoring system based on body weight changes, CD4 counts, hematocrit and clinical concerns identified by the veterinary staff. Euthanasia was performed using an overdose of pentobarbital, while under ketamine anesthesia in accordance with AVMA guidelines, and all efforts were made to minimize suffering.

Plasma samples were obtained during the biannual health screening of macaques at the WaNPRC and tested with the RV1 and RV2-specific Luminex assays for evidence of RV1 and/or RV2 rhadinovirus infections, which are endemic in the colony [65]. Two juvenile *M. nemestrina* M04203 (1.0 years) and K04199 (0.5 years) were determined to be naïve for both macaque RV1 and RV2 rhadinovirus infections by Luminex-based serological and qPCR molecular assays. Twenty adult macaques in concurrent SIV/SHIV vaccine and therapeutic research protocols were identified as possible RFHV+ donors with high titers of anti-RV1 antibodies. Three adult long-term non-progressors in these studies, *M. fascicularis* 01019 and 98039 and *M. nemestrina* A02241, were identified as RFHV+ donor animals as they exhibited high levels of RV1 rhadinovirus DNA in saliva by qPCR.

### Saliva collection

During routine blood draws after ketamine sedation, saliva samples were collected by keeping the macaques in a face-down position allowing the saliva to drip into a petri dish. To obtain saliva for the experimental transmission studies, donor macaques were given 5 mg of pilocarpine 30 minutes before administration of ketamine to induce salivation. The animals were carefully monitored to prevent choking. Saliva was collected and cells were removed from saliva of 98039 and 01019 by centrifugation at 800 × g for 10 min. The supernatant was diluted 1:1 with sterile PBS and filtered through a 0.45 μ. Whatman syringe filter, which is designed for viscous solutions.

### Inoculation of naïve macaques

The saliva preparation from 01019 (3.25 ml) was injected i.v. into M04203 and saliva from 98039 (3.75 ml) was injected i.v. into K04199. Unfiltered whole saliva from A02241 was inserted into the cheek pouches of both M04203 and K04199. Saliva and blood samples were collected weekly and/or bi-weekly from M04203 and K04199. At week 23 post infection, K04199 developed clinical signs meeting the end point criteria for AIDS protocols and was terminated. At weeks 26–29 after the initial saliva infection, M04203 was treated with a regimen of cyclosporine A (15 mgs/kg) IM daily to induce immunosuppression. M04203 was infected with SRV-2B at week 32 and with SIVMne at week 64 to further induce immunosuppression. M04203 was administered dexamethasone 2 mg/Kg daily for 7 days (week 164–165) and then 1 mg/Kg every 3 days from week 166 to week 179 then tapering doses to 0.1 mg/Kg by week 200. M04203 was euthanized at the end of the protocol, week 209. The animals were necropsied and the organs and tissues were examined microscopically.

### SRV-2B isolation and infection

Raji cells were cultured with a frozen buffy coat sample from macaque T81273 diagnosed with KS-like retroperitoneal fibromatosis tumors, obtained from C.-C. Tsai (WaNPRC), and a single infectious clone of SRV-2 (RFS-4) was obtained. Sequence analysis revealed that this SRV-2 clone belonged to the SRV-2B strain associated with the development of retroperitoneal fibromatosis and reactivation of RFHV [74]. At week 32, M04203 was inoculated i.v. with the RFS-4 SRV-2B clone (1 × 10^7^ viral genomes).

### SIVMne infection

Purified preparations of SIVMne that have been used in previous SIV vaccination studies [78, 79] were obtained from S.-L. Hu (WaNPRC). Macaque M04203 was inoculated i.v. at week 64 with SIVMne at 20 50% macaque infectious doses.

### Antibody reagents

Phycoerythrin-labeled anti-human IgG (Jackson ImmunoResearch Laboratories) was used to detect IgG in macaque serum in the Luminex assays.

### Viral load and blood cell quantitation

Blood cell counts and immunophenotyping were performed by the WaNPRC Central Core. PBMC and plasma were separated from blood by gradient centrifugation using Lymphoprep separation medium (Invitrogen). DNA was isolated from PBMC and saliva samples and viral loads were determined using real-time qPCR with TaqMan primers and probes. The SIV qPCR assay [68] and the RV1 [66] and RV2 [34] rhadinovirus assays have been described previously. The SRV-2B qPCR was developed using the forward primer SRV2B-a 5’ CCCGAATTACGATACCAC 3’, the reverse primer SRV2B-b 5’ GTAGCAGTAAGAAGATTAAAGG 3’ and Taqman probe 5’ 56-FAM/TTAGCTTTGCCTAAGGCCCGTG/3BHQ_1 3’ (Integrated DNA Technologies, Inc./ Coralville, IA). The PCR reaction conditions were the same as for the RV2 qPCR and OSM qPCR assays, allowing simultaneous assay runs. The PCR efficiency of the SRV-2B assay was 99.2% (data not shown).

Viral copy number per cell was determined using a qPCR assay targeting oncostatin M (OSM), a single copy cellular gene [66]. The copy number for each assay was calculated from the cycle threshold (C_t_) using the Bio-Rad software. The viral load was determined as a cellular genome copy equivalent using the formula:

Viral load (genome equivalent copies) = Viral copy number/diploid OSM copy number Samples were assayed in duplicate and the means were determined.

### Luminex serological assay

The high throughput multiplex RV1 and RV2 Luminex serological assays were described previously [65]. The EBV Luminex assay was similarly developed using the EBV (B95–8 strain) purified viral lysate (Advanced Biotechnologies, Inc) coupled to Luminex beads. Briefly, the Luminex bead sets coupled to RV1 (KSHV), RV2 (RRV), or EBV virion proteins were incubated with heat-inactivated serum or plasma diluted 1:50 in PBS. Bound antibody was detected using the phycoerythrin-labeled mouse anti-human IgG monoclonal antibody in the Luminex 200HT system (Luminex). The SIV and SRV-2 Luminex assays were performed by the WaNPRC core, as published previously [67].

### Data Analysis

The GraphPad Prism software (San Diego, CA) was used for statistical analysis. For the Luminex data, a cut-off limit of 58 MFI (RV1 assay) and 95 MFI (RV2 assay) was derived from the mean of the non-reactive juvenile sera (RV1 = 9 sera; RV2 = 10 sera) plus 5 standard deviations, as described previously [65].

## Acknowledgments

We acknowledge Patricia Firpo and Shiu-Lok Hu (WaNPRC) for their help in setting up the SIV qPCR assay, LaRene Kueller (WaNPRC) for the SIV and SRV2 Luminex assays, and Courtney Gravett and Elie Karabunarlieva (Rose Lab) for their help in the RV1, RV2 and LCV assays. We also acknowledge Margaret Thouless for the Raji cell culture and advice regarding retrovirus purification and all the staff at the WaNPRC, without whom this research would not have been possible.

